# Lon recognition of the replication initiator DnaA is not confined to a single degron

**DOI:** 10.1101/301655

**Authors:** Jing Liu, Laura Francis, Peter Chien

**Affiliations:** Department of Biochemistry and Molecular Biology, University of Massachusetts Amherst, Massachusetts, 01002; Molecular and Cellular Biology Graduate Program, University of Massachusetts Amherst, Massachusetts, 01002; Department of Biology, University of Massachusetts Amherst, Massachusetts, 01002

**Keywords:** DnaA, degradation, Lon protease, degron, multiple determinants

## Abstract

DnaA initiates chromosome replication in bacteria. In *Caulobacter crescentus*, the Lon protease degrades DnaA to coordinate replication with nutrient availability and to halt the cell cycle during acute stress. Here we characterize the mechanism of DnaA recognition by Lon. We find that the native folded state of DnaA is crucial for its degradation, in contrast to the well-known role of Lon in degrading misfolded proteins. We fail to identify a single degradation motif (degron) sufficient for DnaA degradation, rather we show that both the ATPase domain and a species-specific N-terminal motif are important for productive Lon degradation of DnaA. Mutations in either of these determinants disrupt DnaA degradation *in vitro* and *in vivo.* DnaA switches from an inactive to active state depending on its nucleotide state and we find that locking DnaA in an active state inhibits degradation. Our working model is that Lon engages DnaA through at least two elements, one of which anchors DnaA to Lon and the other acting as an initiation site for degradation.

## INTRODUCTION

DNA replication is a highly regulated process in all life. In bacteria, chromosome replication initiates when the AAA+ protein DnaA assembles at specific motifs in the origin region (1–6). DNA replication occurs only once per cell cycle, and strict regulation of DnaA is needed to coordinate replication with growth and division. It has been shown that DnaA function can be down-regulated through direct inactivation by the DnaA homolog protein Hda and replication clamp, which promotes ATP hydrolysis and convert DnaA from active DnaA•ATP to inactive DnaA•ADP form (7–9). In addition, DnaA activity is also controlled by limiting the interaction between DnaA and the replication origin, which includes the presence of competitive origin binding proteins such as SeqA in *Escherichia coli* (10–12) and CtrA in *Caulobacter crescentus* (13, 14). Finally, DnaA can also be sequestered to non-origin regions of the chromosome as part of its regulation (15, 16). Together, these multiple redundant regulatory mechanisms ensure precise control over replication initiation.

In addition to functional regulation, levels of DnaA are also tightly controlled by expression (17, 18) as well as tuning DnaA lifetime through regulated protein degradation (19, 20). In *Caulobacter crescentus*, DnaA degradation is principally dependent on the Lon protease, while ClpAP plays an additional role in eliminating excess DnaA at specific growth stages (21). Interestingly, Lon-dependent degradation increases when the DnaK chaperone is depleted, which has been attributed to an allosteric stimulation of Lon activity by the excess accumulated misfolded substrates, allowing cells to halt replication during proteotoxic stress (20). Proteolysis also helps rapidly lower DnaA levels upon carbon starvation and nutrient depletion upon entering stationary phase, supporting its important role in sensing environmental changes (22). Interestingly, recent studies found that DnaA degradation is associated with its functional state. Mutations locking DnaA in an ATP-bound form or impairing HdaA-mediated DnaA inactivation led to decreased DnaA stability (21, 23). Although some aspects of recognition motifs for Lon have been described, such as its preference for hydrophobic residues (24), these studies principally center on Lon’s ability to recognize unstructured or poorly folded proteins. Therefore, how Lon recognizes a folded substrate like DnaA remains an outstanding question.

In this study, we address how Lon recognizes DnaA for degradation. Denaturation of DnaA prevents Lon degradation, implying that the native protein fold is important for protease recognition. To investigate which segments of DnaA were important for recognition and degradation of DnaA by Lon, we generated soluble DnaA truncations by limited trypsinization and developed a bioinformatic augmented mass spectrometry strategy to map those fragments. Using this information, we recombinantly produced soluble DnaA fragments, and found that the DnaA AAA+ ATPase domain is sufficient for Lon binding, but not degradation. We also find that a section of the N-terminus specific to *C. crescentus* contributes to degradation. Interestingly, fusing this motif to a normally non-degraded DnaA from another species led to degradation by Caulobacter Lon protease, but this motif does not function as a degron for unrelated proteins. Finally mutations that promote an active conformation of DnaA inhibit Lon recognition. Taken together, our results show that multiple DnaA determinants are needed for Lon recognition with some acting to anchor DnaA to Lon and others acting as initiation sites for degradation. This need for multiple degradation elements likely extends to other protease substrates where single degrons are not easily identified.

## RESULTS

### DnaA needs to be in native folded state to be effectively degraded by Lon protease

The Lon protease is best known as a quality control protease that degrades damaged or misfolded proteins. However, during characterization of DnaA degradation by the Lon protease, we found that denaturing DnaA by heating or urea abolishes its degradation by Lon (Fig. 1A). This suggests that the native protein structure is needed for Lon protease recognition. Note that for these and subsequent reactions, we added the unfolded Lon substrate CMtitin as DnaA is only degraded by Lon *in vitro* when Lon is allosterically activated by an unfolded substrate (20).

**FIGURE 1.**
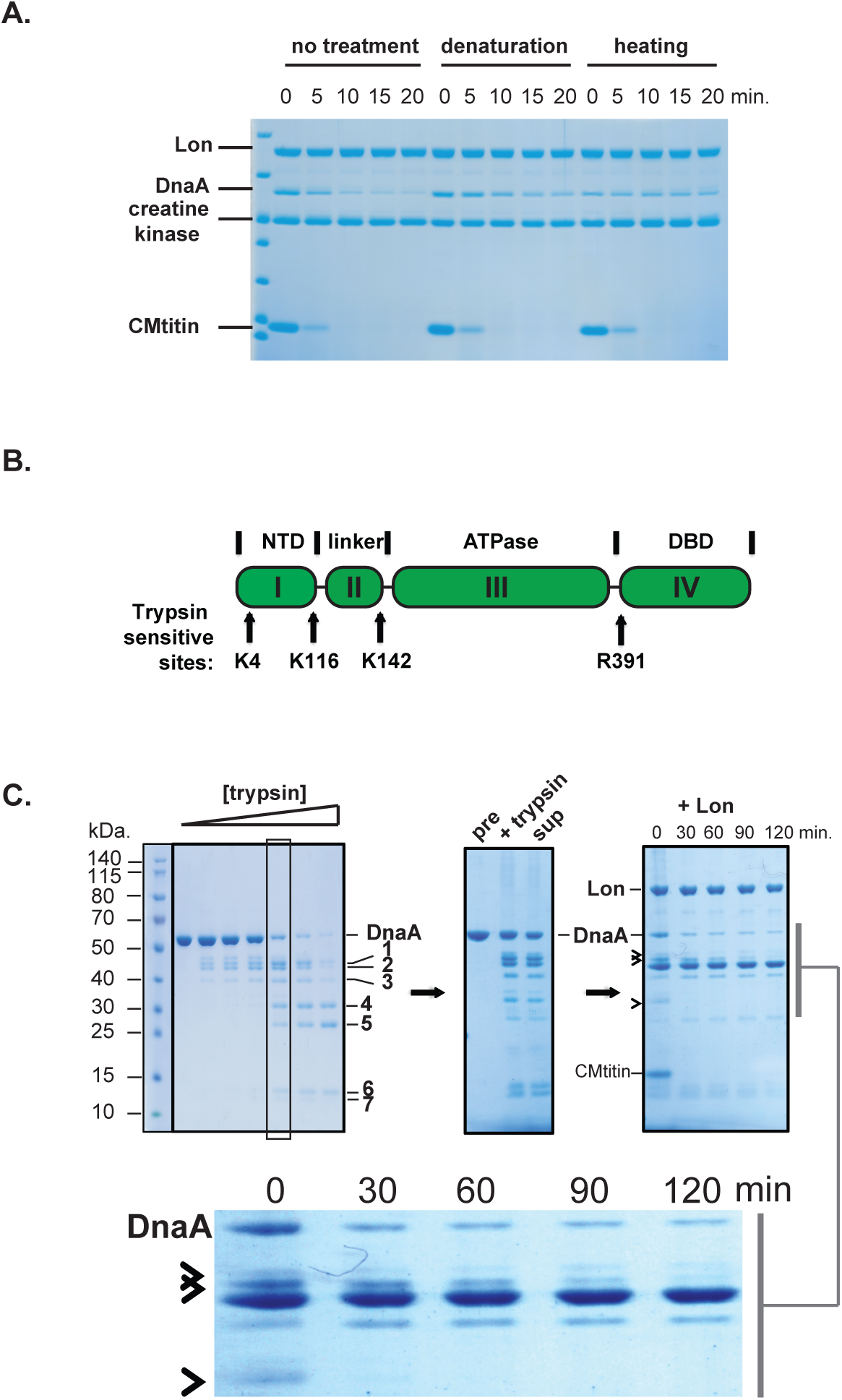
Specific proteolytic determinants reside in different DnaA domains. A. DnaA degradation by Lon was disrupted when DnaA was denatured with 6M urea or heat (45 °C). B. Cartoon of DnaA domains, with predicted limited trypsin sensitive sites marked. C. Limited trypsinization generates several fragments (1–7) that remain in the soluble supernatant upon centrifugation (sup). Some fragments in this mixture (marked with arrowheads) are degraded by Lon (contrast enhanced image shown below).

### Limited trypsination of DnaA identifies regions needed for Lon recognition

DnaA has four conserved functional domains: the N-terminal domain (I), a linker region (II), the AAA ATPase domain (III), and a DNA binding domain (IV) (27) (Figure 1B). To determine the minimal domains in DnaA that were important for recognition and proteolysis by Lon, we first simply assigned termini based on predicted domain boundaries defined by sequence homology to create recombinant fragments of DnaA. Unfortunately, these attempts generally failed to yield soluble proteins, limiting further biochemical characterization (data not shown).

Faced with this problem, we used limited trypsin digestion (which cleaves specifically after Arg/Lys residues) to experimentally determine suitable domain boundaries. Limited proteolysis is a well-established technique that can reveal medium order structure indicated by the accumulation of stable domains during the course of digestion (Figure 1B). We used a range of trypsin concentrations and a fixed time to digest DnaA into several distinct fragments ranging from 10-50 kDa that remained soluble (Figure 1C). To test if any of those fragments were susceptible to Lon-dependent proteolysis, we monitored degradation of the entire fragment mixture in the presence of Lon, ATP, and CMtitin. Several fragments were clearly degraded by Lon while others were stable (Figure 1C). Our interpretation of this result is that the degradable fragments are those that contain the minimal elements needed for Lon engagement.

To map these fragments to the DnaA sequence we first used MALDI mass spectrometry of the trypsination reaction, anticipating that we could uniquely locate the boundaries based on mass. However, the high abundance of Arg/Lys in DnaA resulted in multiple possible fragments that fall into similar mass ranges. Therefore, we developed a bioinformatic program to augment this limited information (see Methods). The prediction program is based on the progressive nature of limited trypsination. First, we assume that under very limiting trypsin conditions, the first fragments derive from only a single cleavage event. Next, we assume that smaller fragments are progressively generated from additional cleavages of the existing larger fragments as more trypsin is used. As a consequence of these assumptions, the true fragment(s) tend to be the one containing re-occurring digestion sites. We then iterate this analysis until we identify unique fragment(s).

With our predictive approach, the DnaA fragments generated by limited trypsination (Figure 1C and Table S2; labeled 1-7) were predicted to arise through cleavage at four internal sites: K116, K142, R391 and K4 (Figure 1B). The first three sites reside in domain boundaries as predicted by sequence homology, thus providing different combinations (domain I, II+III, I+II+III, II+III+IV, III, III+IV and IV). Cleavage at K4 occurs only at higher trypsin concentration and yields domain I without the first four residues at N-terminus. To validate our predictions, we performed Edman degradation on two candidate fragments (labeled 4 and 6 in Figure 1C) and determined that these N-terminal residues matched our prediction (data not shown). Based on this information, we engineered and purified recombinant DnaA proteins containing either native N-/C-termini, or termini corresponding to cleavages at residues K116, K142 and R391.

*In vitro* degradation reactions with the recombinant proteins produced according to our prediction approach showed the same degradation profile as the initial mixture (Figure 2A, 2B comparing with Figure 1C). The summary results of our degradation reactions are: (1) The N-terminal domain I (residues 2-116) is not degraded; (2) The remaining portion of DnaA without the N-terminal domain (117-490) is degraded; (3) The middle linker domain II and ATPase domain III (residues 117-391) is degraded. (4). The C-terminus/domain IV alone (residues 392-490) is not degraded by Lon. Together, the composition of fragments degraded by Lon highlights the importance of the linker region (117-142) as well as the ATPase domain (143-391) for recognition by Lon protease.

**FIGURE 2.**
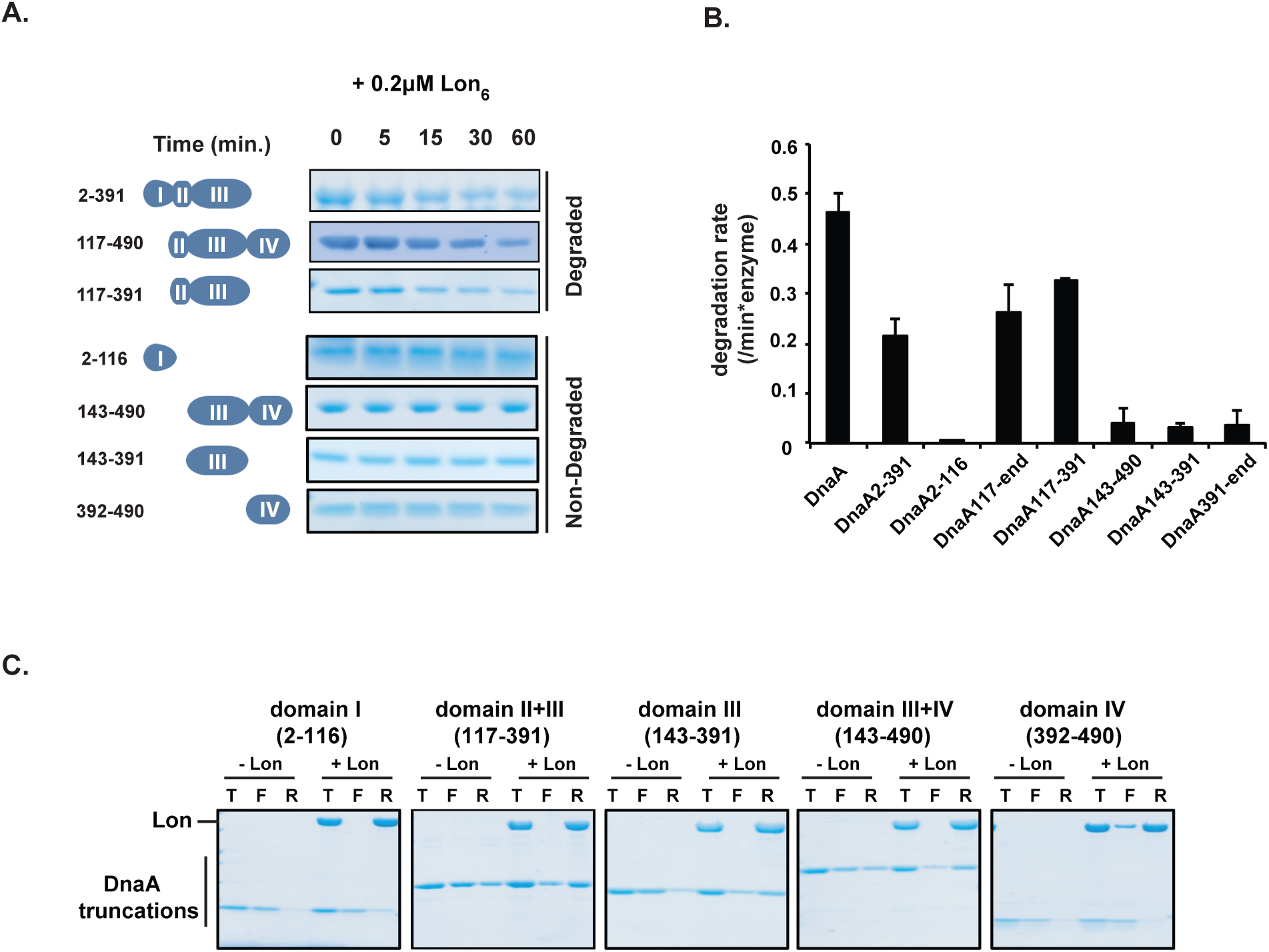
Purified DnaA fragments reveal degradation-prone and Lon-binding regions. A. *In vitro* degradation of purified DnaA domains by the Lon protease. Numbers indicate residue boundaries for each fragment, cartoons of domains shown for each fragment. B. Quantification of DnaA and fragment degradation rates. Means are from three separate experiments for each construct, error bars are standard errors. C. Binding of DnaA truncations and Lon was monitored using spin filtration through a 100kDa cutoff membrane (**T:** total; **F:** filtrate; **R:** retentate) as a proxy for binding. Under these conditions the Lon protein is retained by the membrane, while free DnaA fragments are in the filtrate (Lon; F) and fragments of DnaA that bind Lon are retained (+Lon; R).

*The AAA* + *ATPase domain of DnaA binds Lon.* Next, we tested if these fragments could directly bind Lon. For these experiments, we used an ultrafiltration spin concentrator with a 100-kDa cutoff membrane, which is sufficient to retain the Lon protease, but allows smaller DnaA fragments to flow through. Therefore, DnaA fragments interacting with Lon will be retained by the membrane in the presence of Lon. Using this assay, we found that DnaA fragments corresponding to domains II+III, domain III alone, and domains III+IV bound Lon. By contrast, there was less retention of the isolated domains I and IV by Lon (Figure 2C). These results implicate the AAA+ ATPase domain III of DnaA as necessary for Lon binding. However, domain III alone is not degraded by Lon (Figure 2A).

*Terminal extensions are required for DnaA degradation.* When we compared the degradable and nondegradable DnaA truncations, the domain II linker region (117-142) appeared important for degradation by Lon (Figure 2). For example, Lon degradable DnaA fragments that contain this linker at the N-terminus could be stabilized when this linker is removed (Figure 2; compare 117-391 with 143-391, and 117-490 with 143-490). However, based on the binding data (Figure 2C), this linker region is not important for Lon binding, suggesting it may play a specific role to initiate Lon proteolysis/unfolding.

To test whether this linker specifically played a role in Lon recognition we recoded this linker region using a frameshift strategy. We removed the first nucleotide of codon 117, and appended an additional nucleotide at codon 143, so only the sequence in the linker region was recoded to create the frameshift variant DnaA^*fs*^ (Figure 3A). We purified DnaA^*fs*^ and tested its degradation by Lon. Unexpectedly, this construct was still degraded by Lon (Figure 3B). Moreover degradation of this construct by Lon required the addition of an allosteric Lon activator (CMtitin), which we showed previously to be required for degradation of wildtype DnaA (20). We interpret this to mean that the specific sequence of the linker region is not crucial for degradation of full-length DnaA.

**FIGURE 3.**
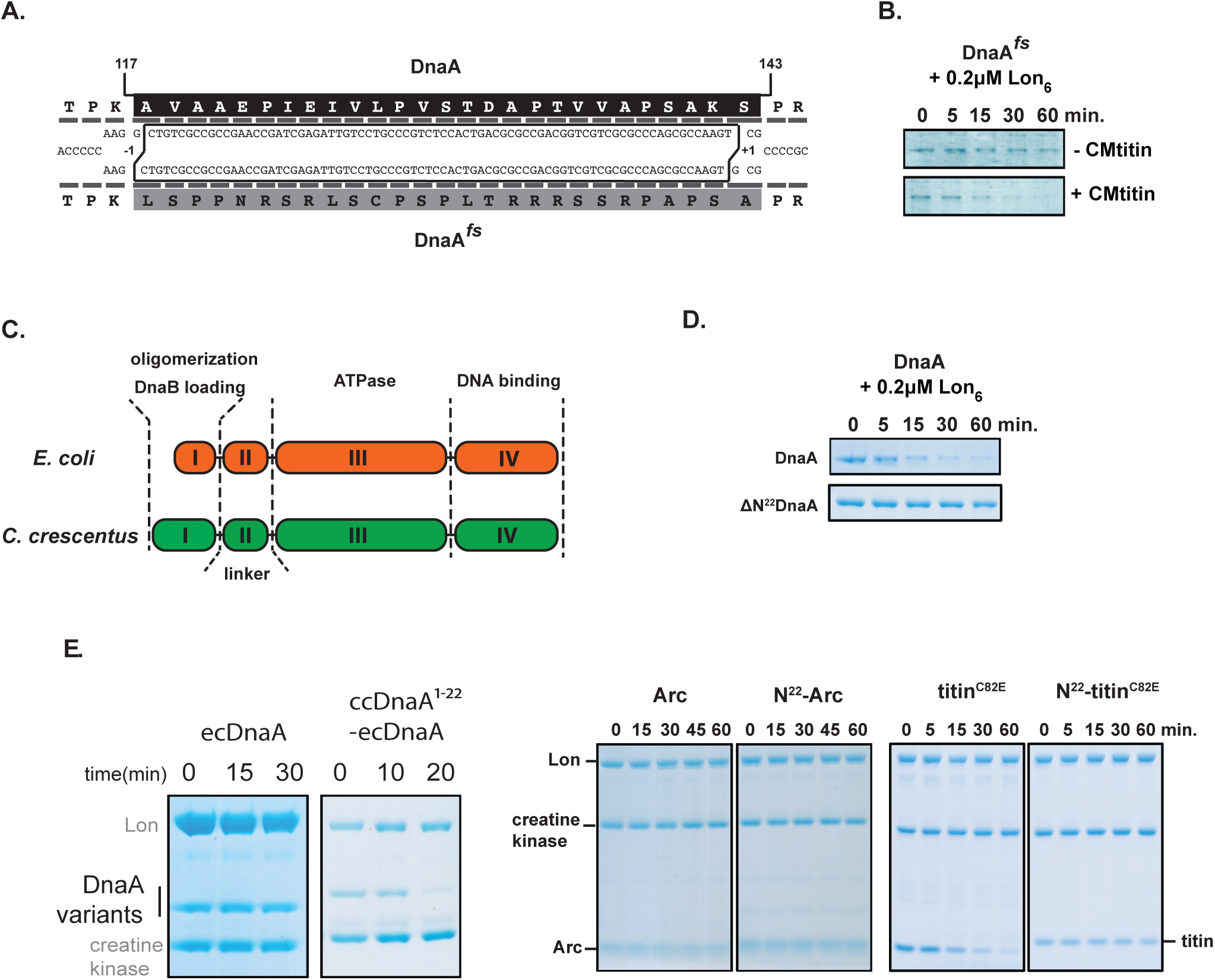
N-terminus extension drives degradation only in the context of DnaA. A. Frame shift mutation in DnaA that recodes sequence in the linker region (117-143). B. Frame shift DnaA mutant is still susceptible degraded by an allosterically activated Lon. Note that Lon in the absence of CMtitin fails to degrade DnaA^fs^, similar to what has been reported for wildtype DnaA (20). C. *Caulobacter* DnaA (which is degraded) has an N-terminal extension compared with *E. coli* DnaA (which is not degraded). D. Degradation of full-length DnaA and DnaA with N-terminal 22 amino acids removed (ΔN^22^DnaA). E. Degradation of *E. coli* DnaA (ecDnaA) or an ecDnaA construct appended with the unique N-terminus of *Caulobacter* DnaA (N22-ecDnaA) by purified *Caulobacter* Lon protease. F. Lon degradation of Arc or titinC82E constructs with or without the additional N-terminal *Caulobacter* DnaA motif, illustrating that the N-terminal motif alone is not sufficient for degradation.

Why does the linker play an important role in degradation of DnaA fragments, but not for the full-length DnaA? One possibility is that there is another element present at the true N-terminus of wildtype DnaA that plays a similar role as the linker with respect to Lon recognition. The *Caulobacter* DnaA contains a longer N-terminus than the *E. coli* DnaA (Figure 3C) an ortholog that is not degraded *in vivo* (40) nor *in vitro* (Figure 3E). To test our hypothesis that the *Caulobacter* N-terminal extension is needed for degradation, we purified a variant of DnaA that lacks this N-terminal extension unique to *Caulobacter* DnaA and found that truncation was resistant to Lon proteolysis (Figure 3D). Interestingly, fusing this extension to the N-terminus of E. coli DnaA resulted in a variant that could now be degraded by Lon (Figure 3E). However, when we fused this motif to other commonly used Lon reporter substrates (Arc and a variant of the titin-I27 domain where a C82E mutation results in destabilization) there was no enhancement of degradation and in the case of titin C82E we found inhibition of degradation (Figure 3F). From these results, we conclude that the N-terminal *Caulobacter* unique extension alone is insufficient for degradation, but that it requires another determinant in DnaA, which likely resides in the AAA+ ATPase domain III and conserved in *E. coli* DnaA, to support Lon degradation.

*Changes in either degradation determinants disrupt DnaA stability in the cell.* Our *in vitro* results indicate a need for the ATPase domain III and N-terminal 22 amino acids of DnaA as Lon recognition elements, and we hypothesized that mutations in these regions would also affect DnaA degradation in the cell. The role of the AAA+ ATPase domain in DnaA degradation has been partly explored in prior work with mutations in the ATPase domain (R357A) resulting in variants of DnaA that are poorly degraded *in vivo* and *in vitro* (21, 23)(Figure 4B). To test the importance of the N-terminal extension, we appended an M2-FLAG epitope tag (DYKDDDK) to the N-terminus of DnaA, resulting in a tagged construct which supports viability (20, 28). Consistent with a role for the specific native N-terminus in DnaA degradation, we found that M2DnaA was resistant to degradation in the cell (Fig. 4A). To further validate this result, we cloned and purified recombinant M2DnaA and found that it also failed to be proteolyzed by Lon *in vitro* (Fig. 4B). Taken with our domain analysis above, our working model is that both the native N-terminus and the ATPase domain III are critical for robust recognition by the Lon protease.

**FIGURE 4.**
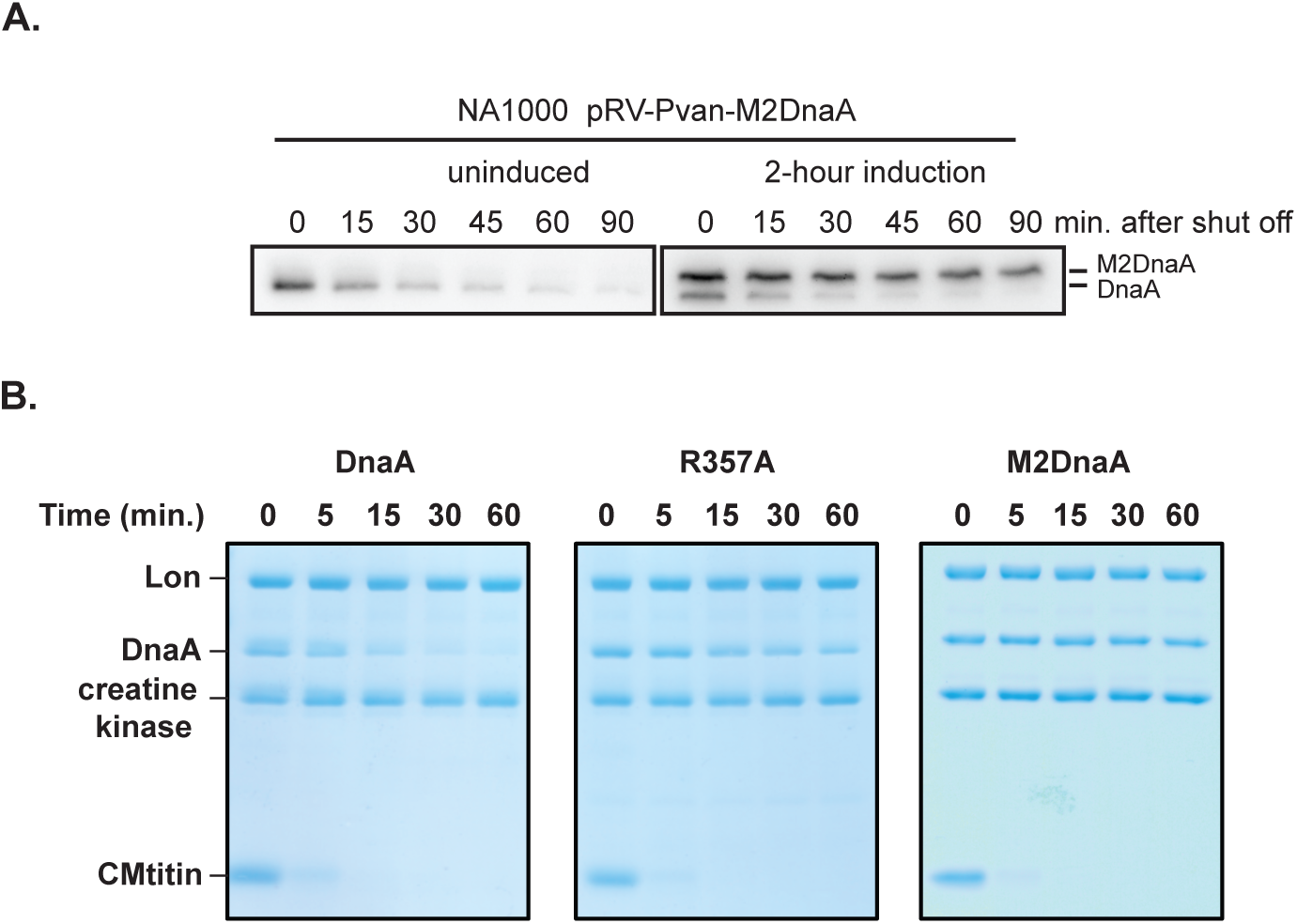
Mutations in domain III and N-terminus produce protease-resistant DnaA variants. A. An M2-FLAG-tagged DnaA (M2-DnaA) is resistant to *in vivo* degradation when expressed in *Caulobacter crescentus.* Degradation was monitored by addition of chloramphenicol to inhibit translation, then levels of DnaA were visualized by Western blots. For the right hand gel, M2-DnaA was induced from a plasmid for two hours prior to translation shut off. B. Purified DnaA mutants with changes in the AAA+ ATPase domain (R357A) or extensions of the N-terminus (M2-DnaA) show reduced degradation compared to wildtype DnaA.

## DISCUSSION

The regulated proteolysis of DnaA by the Lon protease in *Caulobacter crescentus* is an important aspect of the nutrient and proteotoxic stress responses of this bacterium. In our work, we show how different regions of DnaA contribute to post-translational regulation by the Lon protease. We find that the AAA+ ATPase domain of DnaA is critical for Lon binding, providing support for prior observations that the nucleotide-bound state of DnaA affects its stability *in vivo* (23). Our work also suggests that DnaA recognition by Lon occurs through different means than those already characterized for Lon substrates. Namely, Lon is known to degrade misfolded proteins thought to be recognized upon loss of their folded structure, where exposure of hydrophobic patches or stretches of aromatic-rich residues normally buried in the native structure are signals for degradation (24, 32–34). In addition, other substrates have been shown to be recognized by Lon through sequence-based single degrons such as seen with SulA (35), UmuD (36), SoxS (37). Our finding that multiple sequence elements of DnaA are needed for Lon recognition suggests that other Lon substrates may also have multiple elements that must be engaged simultaneously to ensure robust degradation rather than a single constrained degron.

Our working model is that for full-length DnaA degradation, the ATPase domain III binds to Lon and anchors DnaA to the protease, while the N-terminal extension is used as the initiating recognition site for Lon engagement (Figure 5). Alternately, the N-terminal extension could serve as a modulator of the ATPase domain conformation. However, because both removal of the N-terminal extension and masking of the N-terminus through epitope tagging (Figure 4) cause a loss in DnaA degradation, we infer that this latter model is less likely. Our first model of anchoring/initiating is also consistent with our characterization of DnaA fragment degradation. In these experiments, either the natural N-terminus or a linker region at the N-terminus is sufficient to work in concert with the AAA+ ATPase domain III to promote Lon degradation (Figure 2).

**FIGURE 5.**
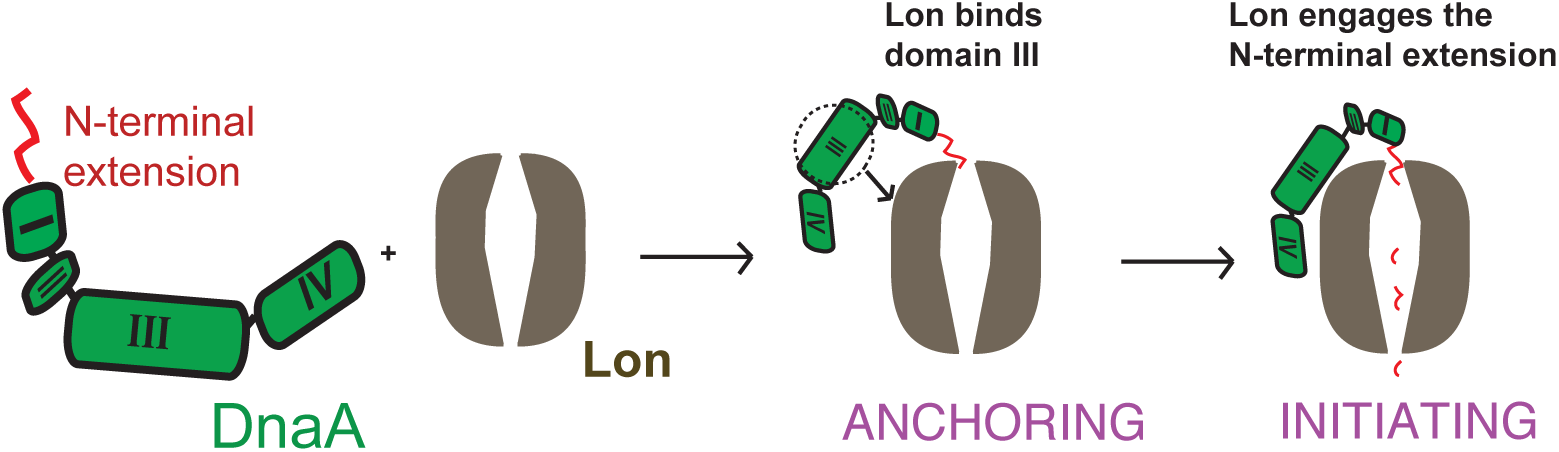
Model of DnaA recognition by Lon protease. Lon degradation of DnaA requires at least two elements. The AAA+ ATPase domain III acts as an anchoring site that binds Lon, but on its own, this is insufficient for degradation. The N-terminal extension of *Caulobacter* DnaA acts as an initiation site for DnaA degradation.

A similar model of anchoring/initiating has been described for other AAA+ proteases. For example, degradation of MuA by the ClpXP protease requires recognition of the MuA complex via “enhancement tags” that promote ClpXP engagement at distal initiation sites more productively (38). Similarly, the 26S proteasome recognizes the structured aspects of ubiquitin, but only initiates proteolysis at an unstructured region in the tagged substrate (39). Our findings reinforce this growing understanding that the degradation signal for a protease may not reside in a single motif and separate anchoring or initiating sites in a given substrate may oversee its destruction.

## EXPERIMENTAL PROCEDURES

*Strains*, *growth conditions and cloning*-A list of strains is shown in Table S1. DnaA truncations and variations were cloned with an N-terminal His_6_-SUMO tag, which could be cleaved during purification by the Ulp1 protease. His_6_-SUMO tagged constructs were ligated into expression vector pBAD33 between unique NdeI and SbfI sites, and transformed into *E. coli* for expression and purification. Arc and its fusion constructs were cloned after His_6_-SUMO in pET23 vector between unique AgeI and NotI sites and transformed into BL21 pLysS strain (Invitrogen). Titin-I27, titin-I27^C82E^ and the fusion construct were cloned after His_6_ tag in pSH21 vector between XbaI and XhoI sites and also transformed into BL21 pLysS strain. *E. coli* expression strains were grown at 37 °C degree. Antibiotics were used when needed with the following concentrations: 30 μg/ml of chloramphenicol and 100 μg/ml of ampicillin. For M2DnaA expression in *Caulobacter crescentus*, the *dnaA* gene containing an extra 5’ CACC was first cloned into pENTR D/TOPO vector (Thermo Fisher Scientific), followed by a recombination with Gateway LR Clonase II enzyme mix (Thermo Fisher Scientific) into a low-copy plasmid destination vector pVan that carries M2-FLAG epitope tag in frame with the downstream insert. This plasmid was transformed into *Caulobacter crescentus* (NA1000) and grown in PYE at 30 °C with kanamycin (5 μg/ml liquid; 10 μg/ml plate) added for plasmid selection.

*Protein purification and modification*–Lon protease was purified as described (24, 25), with additional ion-exchange MonoQ column purification when necessary. The “titin” construct used here is an N-terminally His6 tagged, C-terminally degron β20 tagged I27 domain. The titin^C82E^ is a mutated I27 domain without the C-terminal degron β20. Titin, titin^C82E^, N^22^-titin^C82E^, Arc and N^22^-Arc were purified with Ni-NTA columns as described (24), followed by an additional polishing step with a GE superdex 75 size exclusion chromatography column in H-buffer (25mM HEPES PH7.5, 100mM KCl, 10mM MgCl_2_, 10% Glycerol (v/v) and 1mM DTT). CMtitin was generated by carboxyl methylating two cysteines in titin-I27 with iodoacetamide under urea denaturation condition as described. Modified protein was stored at 4°C in TK buffer (25mM Tris PH8.0, 100mM KCl, 10mM MgCl_2_ and 1mM DTT). DnaA was purified as his_6_SUMO tagged protein and cleaved as described (26), but DnaA purification was carried in S-buffer (20% Sucrose, 25mM HEPES PH7.5, 200mM L-Glutamic acid potassium, 10mM MgCl_2_ and 1mM DTT), and further polished with an additional ion-exchange column (GE healthcare, MonoS G5/50) using a KCl gradient from 0.1M to 1M in MS-S buffer (20% Sucrose, 25mM Tris PH8.5, 2mM DTT).

*Limited trypsinization, Lon*-*dependent degradation of fragments and MALDI mass spectrometry*-To perform limited trypsin digestion, a serial dilution of trypsin starting with 10μg/ml were added to 10μM DnaA in S-buffer and incubated at 25°C for 30 minutes. 5 mM protease inhibitor phenylmethylsulfonyl fluoride (PMSF) was added to quench the reactions. Part of the resulting digest was used for SDS-PAGE analysis, part for MALDI, and the rest was used for the Lon degradation assay. To perform Lon degradation on digested fragments, the reactions were buffer-exchanged into fresh S-buffer (no PMSF) with polyacrylamide spin desalting columns (ThermoFisher Scientific) and the additional components (Lon and ATP regeneration mix) were added immediately after the spin to initiate the reaction. For MALDI mass spectrometry, the digested fragments were precipitated by TCA and resuspended in 5% formic acid. MALDI mass spectrometry was performed by the UMass Amherst / IALS Mass Spectrometry Center.

*Bioinformatic prediction of trypsinized fragments*-Our algorithm to predict cleavage profiles is based on the most likely initial cleavage sites, as well as cleavage re-occurrence. The first iteration of cleavage prediction identifies several combinations of cleavage sites with different probabilities of being correct and we assign each site a confidence score. Next, we scanned through the experimental dataset to find fragments with at least one highest scored cleavage sites, and increased the confidence score of the other end of the fragment if its score was lower, as we assume that this fragment is a product of further digestion that was not found on the initial run. This process was repeated multiple times, until all the most likely fragments harboring highest confidence scores at both ends of a predicted fragment emerges from the dataset. We implemented the prediction algorithm in Python (available upon request). To ensure we include all possible digestion pairs, we assigned a high tolerance range to ±1% precision on each MALDI determined masses (e.g. the determined mass could be off by 300 Da from its real mass for a 30kDa protein, Table 1). After multiple iterations of scoring and refinement, we were able to converge on one specific Arg/Lys pair for each major fragment.

*In vitro degradation assay*–Degradation for all constructs were performed at 30°C with following protease concentration unless elsewhere indicated: 0.2μM Lon_6_, 1.5 μM DnaA or ΔN^22^DnaA, 2 μM other truncation fragments expect 2-116 (5 μM) or 391-490 (10 μM), with 4 mM ATP, 15 mM creatine phosphate (Sigma) and 75 μg/ml creatine kinase (Roche) as ATP regeneration components. The reaction is initiated by adding ATP regeneration mix to the protease-substrate solution in TK buffer (25mM Tris PH8.0, 100mM KCl, 10mM MgCl_2_ and 1mM dithiothreitol, and the reaction was carried at 30°C. 10μl aliquots were taken at each time point and quenched with SDS loading dye (2% SDS, 6% Glycerol, 50mM Tris PH8.0 and 2mM DTT), and examined by SDS-PAGE. Using creatine kinase as an internal loading control, the degradation rate was determined by protein band intensity change at different time points analyzed with ImageJ 1.47(NIH) software.

*Filtration*-*spin Lon binding assay*-To set up the assay, 120 μl reactions containing different DnaA domains were incubated with or without Lon protease at 30°C for 10 minutes with following concentrations: 5 μM DnaA truncations, 0.5 μM Lon_6_ and 1mM ATP-γ-S in TK buffer. 20μl of the mixtures were taken as the control for total input, and the remaining samples were transferred to a Vivaspin 500 concentrator (100kDa, Viva Product) and spun down at 15,000 xg for 10 minutes. Flowthrough fraction was collected, and the column was washed with 120 μl TK buffer containing 1mM ATP-γ-S and 0.05% Tween-20. Proteins in the retention fraction were re-suspended in 120μl TK buffer. SDS loading dye was added to input, flow-through and retention fractions. Samples were heated at 95°C for 10 minutes and analyzed by SDS-PAGE/Coomassie staining.

## Acknowledgments

We thank the Sauer lab (MIT) for sharing the pSH21-titinI27-β20 plasmid to generate titin-I27 substrate and related proteins, Stephen Eyles (UMass Amherst) for mass spectrometry experiments and discussions. Hanwei Zhao from Kaltashov lab (UMass Amherst) and Xiaojian Wu from Zilberstein lab (Computer Science department, UMass Amherst) for discussions on bioinformatic predictions on limited proteolysis. We thank members of the Chien lab, the Protein Homeostasis theme of the Center for Models to Medicine, and Sloan Siegrist for helpful feedback. This work was sponsored in part by funding from a Chemistry Biology Interface Program Training Grant (NIH T32GM08515) to J.L., and from National Institute of Health Grant R01GM111706 to P.C.

## Conflict of interest

The authors declare that they have no conflicts of interest with the contents of this article.

## Author contributions

P.C. and J.L. conceived the idea for the project. J.L. performed experiments, developed new reagents, and analyzed the results. L.F. purified DnaA and performed experiments. J.L. and P.C. wrote the paper.

## Supplemental information

**Table S1.**

| Strain | Description | Source |  |
| --- | --- | --- | --- |
| ER2566 | general cloning strain | NEB | - |
| BL21 (DE3) | general cloning strain | Invitrogen | Chloramphenicol |
| Acela | general cloning strain | EdgeBio | Chloramphenicol |
| Top10 | general cloning strain | Invitrogen | - |
| EPC481 | BL21 pSH21-Titin-I27B20 | Gur and Sauer. 2008 | Ampicillin |
| EPC517 | BL21 pET23b-hissumo-DnaA | Jonas et al. 2013 | Ampicillin |
| EPC523 | BL21 pET23b-hissumo-delNDnaA | this work | Ampicillin |
| EPC460 | ER2566 pBAD ccLon | Gora et al. 2013 | Chloramphenicol |
| EPC604 | ER2566 pBAD ecLon | this work | Chloramphenicol |
| EPC830 | BL21 pET23DnaA1-22titinC82E | this work | Ampicillin |
| EPC849 | ER2566 pBAD33-hissumo M2DnaA | this work | Chloramphenicol |
| EPC857 | BL21 pET23-HisC82E no tag | this work | Ampicillin |
| EPC878 | Acela pET23-DnaA22-titinC82E | this work | Chloramphenicol |
| EPC1025 | ER2566 pBAD33-hissumo-DnaA2-391 | this work | Chloramphenicol |
| EPC1026 | ER2566 pBAD33-hissumo-DnaA2-116 | this work | Chloramphenicol |
| EPC1027 | ER2566 pBAD33-hissumo-DnaA117-490 | this work | Chloramphenicol |
| EPC1028 | ER2566 pBAD33-hissumo-DnaA117-391 | this work | Chloramphenicol |
| EPC1029 | ER2566 pBAD33-hissumo-DnaA143-490 | this work | Chloramphenicol |
| EPC1030 | ER2566 pBAD33-hissumo-DnaA143-391 | this work | Chloramphenicol |
| EPC1152 | BL21 pET23b-Arcst11 | this work | Ampicillin |
| EPC1153 | BL21 pET23b-N22-Arcst11 | this work | Ampicillin |
| EPC1154 | ER2566 pBAD33-hissumo-ecDnaA | this work | Chloramphenicol |
| EPC1155 | ER2566 pBAD33-hissumo-chimeric ecDnaA | this work | Chloramphenicol |
| EPC1156 | ER2566 pBAD33-hissumo-DnaA 392-490 | this work | Chloramphenicol |
| EPC1157 | ER2566 pBAD33-hissumo-DnaA fs | this work | Chloramphenicol |

**Table S2.**
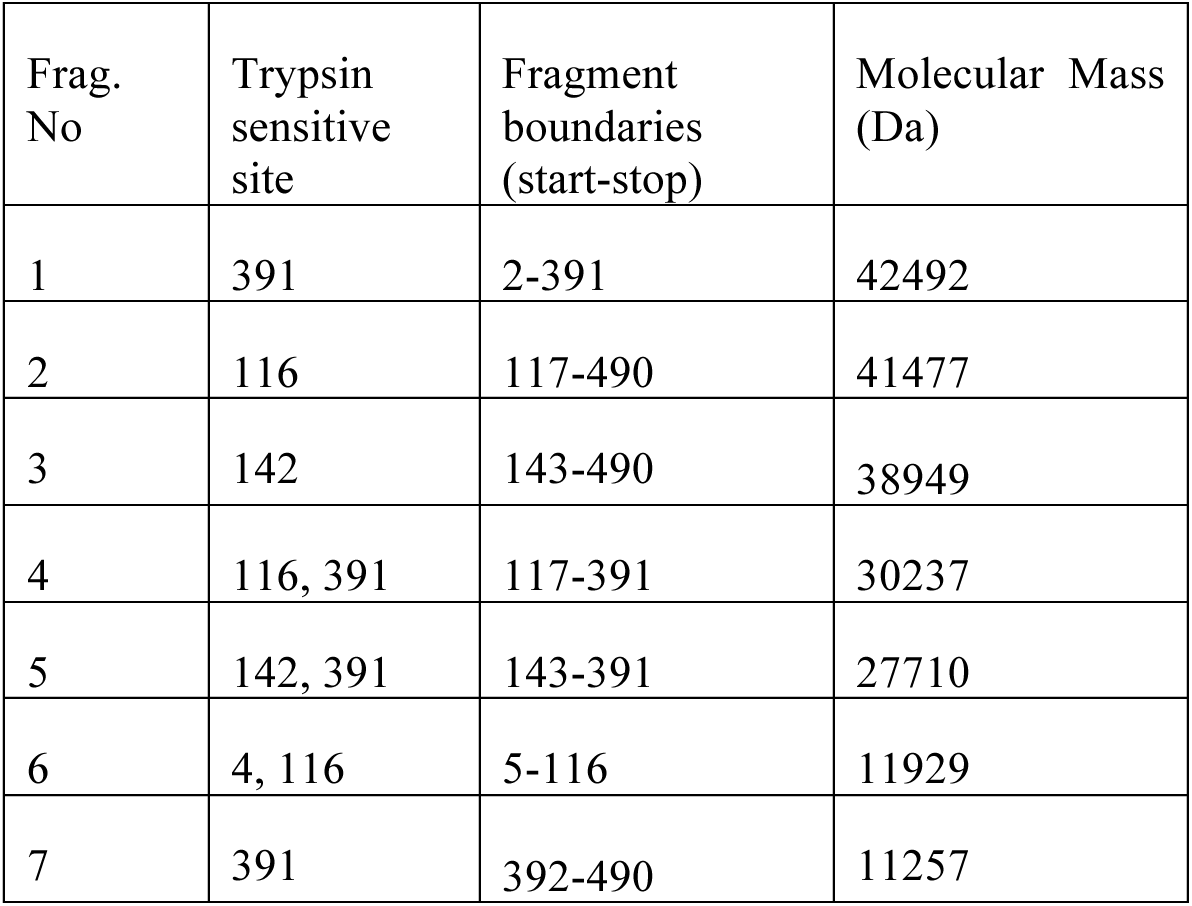
Predicted trypsin digestion fragments (the last residue of DnaA is 490)

| Frag. No | Trypsin sensitive site | Fragment boundaries (start-stop) | Molecular Mass (Da) |
| --- | --- | --- | --- |
| 1 | 391 | 2-391 | 42492 |
| 2 | 116 | 117-490 | 41477 |
| 3 | 142 | 143-490 | 38949 |
| 4 | 116, 391 | 117-391 | 30237 |
| 5 | 142, 391 | 143-391 | 27710 |
| 6 | 4, 116 | 5-116 | 11929 |
| 7 | 391 | 392-490 | 11257 |

